# Antidepressants stimulate lipoprotein(a) macropinocytosis via serotonin-enhanced cell surface binding

**DOI:** 10.1101/2021.03.26.437114

**Authors:** Nikita Deo, Halima Siddiqui, Golnoush Madani, Malcolm Rutledge, Michael JA Williams, Sally PA McCormick, Gregory MI Redpath

**Affiliations:** Department of Biochemistry, School of Biomedical Sciences, University of Otago, Dunedin, New Zealand; HeartOtago, University of Otago, Dunedin, New Zealand; Department of Medicine, Otago Medical School, University of Otago, Dunedin, New Zealand; EMBL Australia Node in Single Molecule Science, School of Biomedical Sciences, University of New South Wales, Sydney, Australia

**Author notes:** ***For correspondence:*** (SPAM), (GMIR).

**Keywords:** lipoprotein(a), macropinocytosis, selective serotonin reuptake inhibitors, PlgRKT, serotonin

## Abstract

We recently found that plasminogen receptors regulate the plasma membrane binding and uptake of Lp(a) via macropinocytosis. In this study, we sought to further define lipoprotein(a) [Lp(a)] macropinocytosis, discovering an unexpected role for antidepressants and serotonin in the regulation of this process. We found that the tricyclic antidepressant imipramine enhanced Lp(a) uptake, in contradiction of its published role as a macropinocytosis inhibitor. Extending these experiments to the commonly used serotonin uptake inhibitors (SSRIs) citalopram and sertraline, we found that citalopram stimulated, while sertraline inhibited, Lp(a) uptake. Imipramine and citalopram enhanced cell surface binding of Lp(a) rather than upregulating macropinocytosis. Consistent with imipramine and citalopram boosting extracellular serotonin levels, serotonin itself also enhanced Lp(a) surface binding and uptake. Imipramine and serotonin increased expression of the plasminogen receptor with a C-terminal lysine (PlgRKT), a receptor known to enhance cell surface binding of Lp(a), likely accounting for their effects on Lp(a) uptake. Finally, imipramine and citalopram increased Lp(a) delivery into Rab11 recycling endosomes, but not degradative pathways in the cell. These findings indicate citalopram and imipramine may have utility as a potential Lp(a)-lowering therapeutic in people suffering from depression who often have elevated Lp(a) levels and an increased risk of cardiovascular disease.

## Introduction

Lipoprotein(a) (Lp(a)) is a highly atherogenic form of low-density lipoprotein (LDL) (Boerwinkle et al., 1992). Lp(a) consists of an LDL-like particle containing apolipoprotein-B (apoB), covalently bound to apolipoprotein(a) (apo(a)), a glycoprotein that is structurally similar to plasminogen (Lawn et al., 1997). Lp(a) is elevated (greater than 100 nmol) in at least 20% of the European population (Wilson et al., 2019) and recent data from the United Kingdom Biobank indicates that every 50 nmol increment increase in Lp(a) imparts a greater risk of developing cardiovascular disease (CVD) (Patel et al., 2020). Elevated Lp(a) levels are frequently overlooked despite having an equal impact on CVD development and risk as LDL (Langsted and Nordestgaard, 2020). Unlike LDL therapies, specific Lp(a)-lowering treatments remain limited, with RNA antisense therapy targeted to apo(a) being the only treatment close to becoming clinically available (Lippi et al., 2020). Understanding how LDL is cleared from circulation by endocytosis has led to the development of statins and newer therapies such as PCSK9 inhibitors (Chaudhary et al., 2017) to lower LDL. Despite the importance of endocytic processes in lipoprotein clearance, little is known about the endocytic regulation of Lp(a). The inability to ascribe a clear single receptor that mediates Lp(a) uptake has hampered the development of effective therapies for Lp(a) lowering.

Multiple receptors have been implicated in binding Lp(a) and mediating Lp(a) endocytic uptake into the cell including the LDL receptor (Floren et al., 1981; Hofmann et al., 1990; Romagnuolo et al., 2017; Tam et al., 1996), VLDL receptor (Argraves et al., 1997) and plasminogen receptors (Sharma et al., 2017), yet in each case the impact of each receptor on Lp(a) uptake is either inconsistent or only partially responsible for uptake (Chemello et al., 2020; McCormick and Schneider, 2019; Rader et al., 1995; Reblin et al., 1997; Romagnuolo et al., 2017), leaving our understanding of the process of Lp(a) catabolism incomplete. Recently, we found that multiple plasminogen receptors regulate Lp(a) uptake via the endocytic process of macropinocytosis (Siddiqui et al., 2023).

There are many pathways of endocytosis identified to date, each with specific cargoes, activation conditions and functions (Redpath et al., 2020). Clathrin-dependent and fast endophilin mediated endocytosis represent mechanisms of receptor-mediated endocytosis. In both cases, cargo/ligand binding to the receptor leads to the assembly of adaptor and coat proteins around the forming endosome which promote receptor and cargo internalisation (Boucrot et al., 2015; Casamento and Boucrot, 2020; Haucke, 2006; Traub and Bonifacino, 2013). In contrast, macropinocytosis is an endocytic pathway where cargo is internalised *via* membrane remodelling events as opposed to receptor-mediated initiation of endocytosis.

Here, we set out to further define Lp(a) macropinocytosis. Using the tricyclic antidepressant imipramine, a recently identified macropinocytosis inhibitor (Lin et al., 2018), we surprisingly found that imipramine *stimulated* Lp(a) macropinocytosis in liver cells. As imipramine partially exerts its antidepressant effect via inhibition of serotonin reuptake, we next sought to determine the effects of commonly used selective serotonin reuptake inhibitor (SSRI) antidepressants on Lp(a) uptake. Citalopram also robustly stimulated Lp(a) uptake in liver cells while sertraline inhibited Lp(a) uptake, likely due to its effects as a dynamin inhibitor. Given the incongruity of the published role of imipramine as a macropinocytosis inhibitor (Lin et al., 2018) and our results, we next defined how imipramine and citalopram regulated Lp(a) uptake. Neither drug affected macropinocytosis directly but rather enhanced Lp(a) binding to the cell surface. Serotonin also enhanced Lp(a) cell surface binding, indicating that imipramine and citalopram may exert their effects by increasing extracellular serotonin levels through serotonin transporter inhibition. We then tested the effects of imipramine, citalopram and serotonin on expression of plasminogen receptors important in Lp(a) uptake (Siddiqui et al., 2023), finding serotonin and imipramine increased expression of the membrane anchor PlgRKT, and serotonin and citalopram increased expression of the macropinocytosis regulator S100A10. Finally, we showed that imipramine and citalopram increased Lp(a) delivery into recycling endosomes, but not late endosomes or lysosomes. These findings establish that serotonin and antidepressants are important regulators of Lp(a) cell surface binding and uptake, which given their clinical importance has potential implications for the treatment of depression and cardiovascular disease.

## Results

### Antidepressants are modulators of Lp(a) uptake

In an effort to further refine how macropinocytosis regulates Lp(a) uptake, we employed imipramine, a tricyclic antidepressant that was recently identified as a macropinocytosis inhibitor (Lin et al., 2018). Unexpectedly, imipramine treatment stimulated Lp(a) uptake in HepG2 cells in a concentration-dependent manner (∼290% increase at 20 μM, Figure 1A-C). Given our unexpected finding, we tested the effects of the more commonly used selective serotonin reuptake inhibitors (SSRIs) on Lp(a) uptake. As with imipramine, overnight treatment with the SSRI citalopram induced robust, concentration-dependent increases in Lp(a) uptake (∼300% increase at 50 μM, Figure 1D-F). In contrast, sertraline had no significant effect (Figure 1G-I). As we have shown Lp(a) uptake occurs via macropinocytosis (Siddiqui et al., 2023), we employed the commonly used marker of macropinocytosis, 70 kDa dextran (Li et al., 2015), to probe the mechanism of antidepressant action. Unexpectedly, dextran uptake was not significantly increased with imipramine or citalopram treatment (Figure S1A,B). In contrast, sertraline had a significant inhibitory effect on dextran uptake (Figure S1A,C), indicating it acts as a macropinocytosis inhibitor. Imipramine and citalopram therefore stimulate Lp(a) uptake independent of macropinocytosis in liver cells, while sertraline may be incapable of enhancing Lp(a) uptake as it concurrently inhibits macropinocytosis.

**Figure 1:**
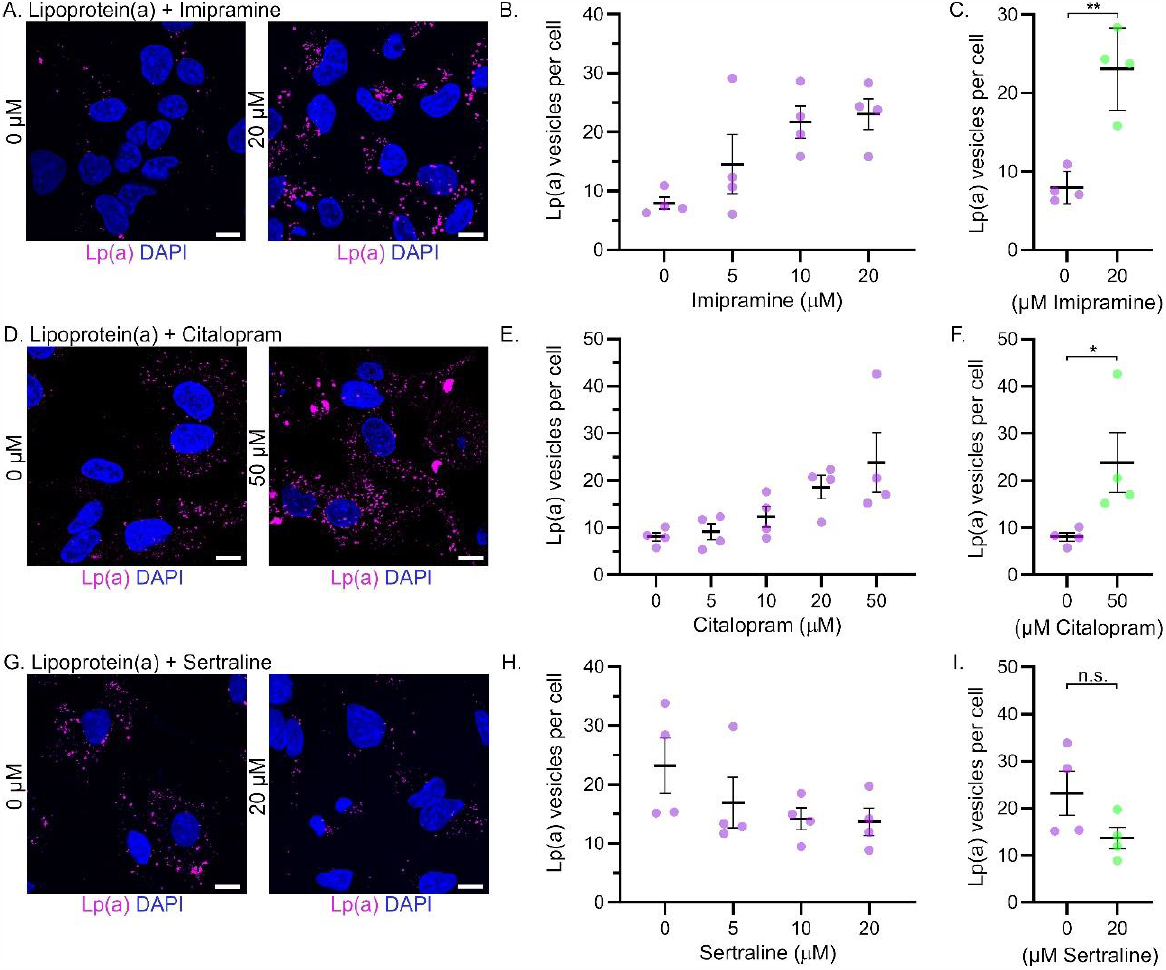
Imipramine and citalopram, but not sertraline, enhance Lp(a) uptake in liver cells. **(A)** Representative images of HepG2 cells incubated with Lp(a) for 1 hour following H_2_O vehicle (left) or 20 μM imipramine (right) treatment overnight. **(B)** Lp(a) vesicles detected following overnight treatment with 5, 10, and 20 μM imipramine compared to vehicle control. **(C)** Lp(a) vesicles detected following overnight treatment with 20 μM imipramine compared to vehicle control. **(D)** Representative images of HepG2 cells incubated with Lp(a) for 1 hour following H_2_O vehicle (left) or 50 μM citalopram (right) treatment overnight. **(E)** Lp(a) vesicles detected following overnight treatment with 5, 10, 20 and 50 μM citalopram compared to vehicle control. **(F)** Lp(a) vesicles detected following overnight treatment with 50 μM citalopram compared to vehicle control. **(G)** Representative images of HepG2 cells incubated with Lp(a) for 1 hour following H_2_O vehicle (left) or 20 μM sertraline (right) for 2 hours. **(H)** Lp(a) vesicles detected following 2 hours treatment with 5, 10, and 20 μM sertraline compared to vehicle control. **(I)** Lp(a) vesicles detected following 2-hour treatment with 20 μM sertraline compared to vehicle control. n.s. = not significant, *= p<0.05, **=p<0.01 from Student’s T-test. Data points represent means of independent experiments quantified from 5 fields of view of ∼20 cells per field. Error bars represent standard error of the mean. Lp(a) was detected using the LPA4 primary antibody and AlexaFluor conjugated anti-mouse secondary antibody. Images were acquired on an Olympus FV1000/FV1200 confocal microscope. Scale bar= 5 μM.

### Imipramine and citalopram stimulate Lp(a) binding to the cell surface

We next sought to understand how imipramine and citalopram enhanced Lp(a) uptake in liver cells. As Lp(a) anchoring to the cell surface is required for its subsequent macropinocytosis (Siddiqui et al., 2023), we hypothesised that imipramine and citalopram would increase Lp(a) binding to the plasma membrane. To test this, we performed cell surface binding experiments. HepG2 cells were treated with imipramine or citalopram overnight, cooled to prevent endocytic uptake, incubated with Lp(a) or dextran, fixed, washed and Lp(a) or dextran-TRITC bound to the cell surface quantified. Consistent with our hypothesis, Lp(a) levels on the cell surface were increased following imipramine and citalopram treatment in HepG2 cells (Figure 2A,B; ∼270% increase for imipramine, ∼190% increase for citalopram). Conversely, dextran surface binding was unaltered by imipramine and citalopram treatment (Figure 2C,D), confirming that the increased Lp(a) macropinocytosis induced by imipramine and citalopram is due to enhanced Lp(a) binding to the plasma membrane.

**Figure 2:**
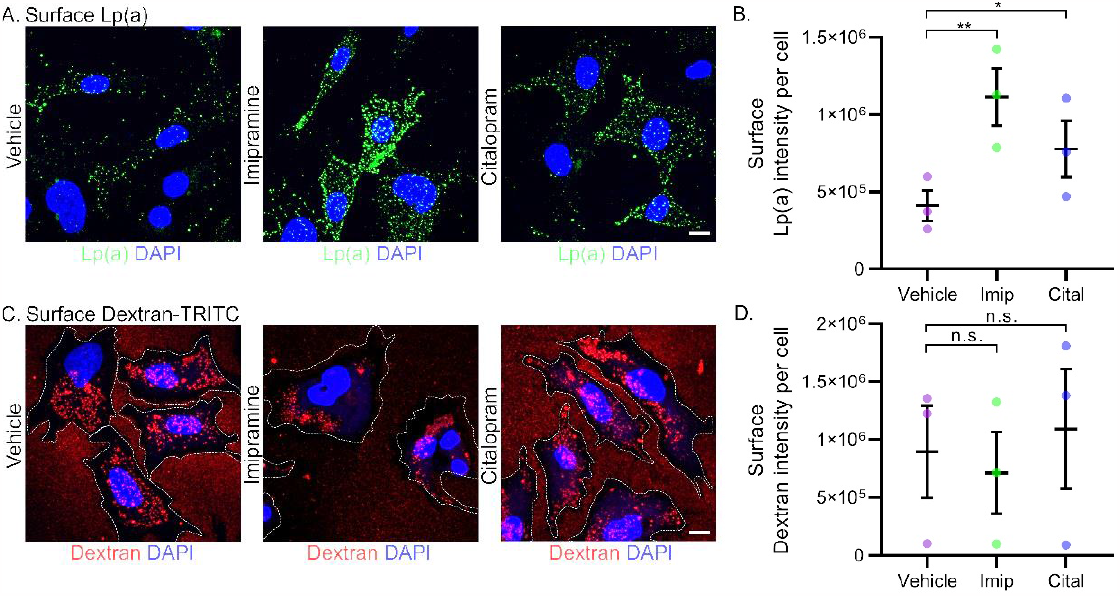
Imipramine and citalopram increase Lp(a) cell surface binding. **(A)** Representative images of maximum z-projections of surface-bound Lp(a) following H_2_O vehicle, 20 μM imipramine or 50 μM citalopram treatment overnight. **(B)** Surface Lp(a) signal per cell in vehicle, imipramine and citalopram treated cells. **(C)** Representative images of maximum z-projections of surface-bound dextran following H_2_O vehicle, 20 μM imipramine or 50 μM citalopram treatment overnight. **(D)** Surface dextran signal per cell in vehicle, imipramine and citalopram treated cells. n.s. = not significant, *= p<0.05, p=<0.01 from randomised-block ANOVA (standard-**A**, Friedman-**B**) with Dunn’s correction for multiple comparisons. Data points represent means of independent experiments quantified from 5 fields of view of ∼20 cells per field from each experiment. Error bars represent standard error of the mean. 70 kDa dextran-TRITC was detected directly. Lp(a) was detected using the LPA4 primary antibody and AlexaFluor conjugated anti-mouse secondary antibody. Images were acquired on an Olympus FV1000/FV1200 confocal microscope. Scale bar= 5 μM.

Since apo(a) is the predominant mediator of Lp(a) uptake (Siddiqui et al., 2023), we repeated surface binding and uptake results using apo(a)-mScarlet to confirm that imipramine and citalopram stimulation of Lp(a) uptake is mediated by apo(a). While imipramine and citalopram induced increases in apo(a) surface binding to HepG2 cells in each of 4 independent experiments (mean surface intensity of 461850 for imipramine, 432856 for citalopram vs. 174446 for vehicle control), this difference did not reach statistical significance (Figure S2A,B). However, imipramine and citalopram induced robust, statistically significant increases in apo(a)-mScarlet uptake into HepG2 cells (∼197% increase for imipramine and citalopram, Figure S2C-E). Together, these results indicate apo(a) mediates the enhanced plasma membrane binding and subsequent uptake of Lp(a) in response to imipramine and citalopram treatments.

### Serotonin enhances Lp(a) cell surface binding and macropinocytosis

The primary target of antidepressants is the serotonin transporter SLC6A4. Serotonin transporter inhibition prevents serotonin uptake from the extracellular space (Andersen et al., 2009), leading to elevated extracellular serotonin levels. Interestingly, a recent study found addition of serotonin to the media cells enhanced transferrin cell surface binding and endocytosis by modulating the biophysical properties of the plasma membrane (Dey et al., 2021). HepG2 cells express the highest level of the predominant serotonin transporter *SLC6A4* of any human cell line in the Human Protein Atlas (Thul et al., 2017), indicating they may be highly sensitive to serotonin transporter inhibition. Given the above, we reasoned that imipramine and citalopram may be predominantly acting by inhibiting the serotonin transporter, leading to increases in extracellular serotonin levels, which could enhance Lp(a) surface binding and uptake similar to that seen for endocytic cargoes in Dey et al., (2021).to increases in extracellular serotonin levels, which could enhance Lp(a) surface binding and uptake similar to that seen for endocytic cargoes in Dey et al., (2021).

To determine if serotonin itself enhanced Lp(a) surface binding and uptake, we treated HepG2 cells overnight with serotonin using the concentrations established by Dey et al., to enhance transferrin surface binding (Dey et al., 2021). We then incubated the cells with Lp(a) on ice to determine surface binding as above. Treatment with 500 μM serotonin resulted in a significant increase (∼230%) in Lp(a) binding to the cell surface (Figure 3A,B) and a comparable increase in Lp(a) uptake (∼260%, Figure 3C-D). Similar to imipramine and citalopram, treatment of HepG2 cells with serotonin resulted in no significant changes in dextran binding to the cell surface (Figure 3E,F) or dextran uptake (Figure 3G,H). These data are consistent with imipramine and citalopram acting by increasing extracellular serotonin levels.

**Figure 3:**
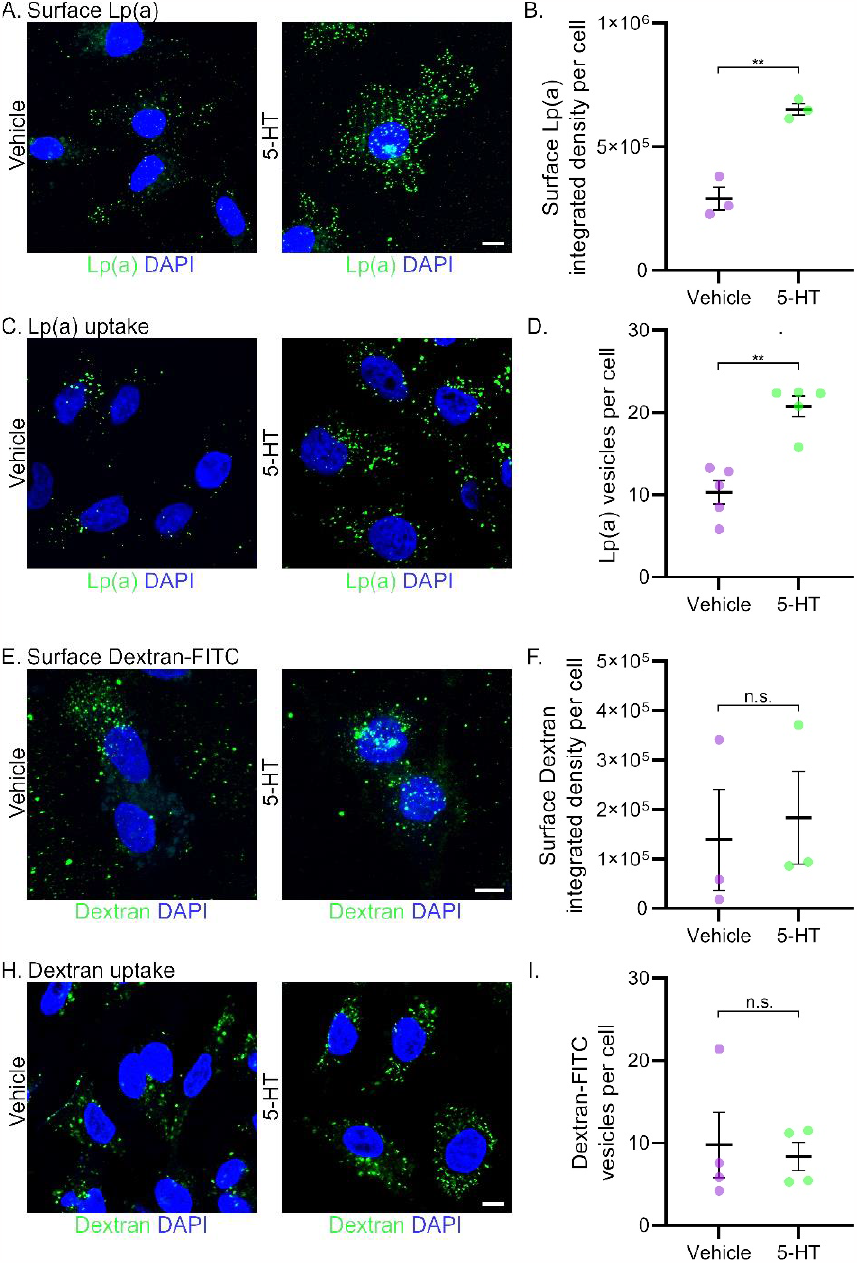
Serotonin increases Lp(a) surface binding and uptake. **(A)** Representative images of maximum z-projections of surface-bound Lp(a) following H_2_O vehicle (left) or 500 μM serotonin treatment (right) of HepG2 cells overnight. **(B)** Quantification of surface Lp(a) signal per cell in vehicle or serotonin treated cells. **(C)** Representative images of endocytosed Lp(a) after overnight treatment of HepG2 cells with H_2_O (left) or 500 μM serotonin (right) and incubation with Lp(a) for 1 hour. **(D)** Quantification of Lp(a) vesicles per cell in vehicle and serotonin treated cells. **(E)** Representative images of maximum z-projections of surface-bound dextran following H_2_O vehicle (left) or 500 μM serotonin treatment (right) of HepG2 cells overnight. **(F)** Quantification of surface dextran signal per cell in vehicle or serotonin treated cells. **(G)** Representative images of endocytosed dextran after overnight treatment of HepG2 cells with H_2_O (left) or 500 μM serotonin (right) and incubation with dextran for 1 hour. **(H)** Quantification of dextran vesicles per cell in vehicle and serotonin treated cells. n.s.= not significant, *= p<0.05, **p=<0.01 from unpaired Student’s T-test **(B)** or Mann-Whitney test **(D**,**F**,**H)**. Data points represent means of independent experiments quantified from 5 fields of view of ∼20 cells per field from each independent experiment. Error bars represent standard error of the mean. 70 kDa dextran-TRITC was detected directly. Lp(a) was detected using the LPA4 primary antibody and AlexaFluor^488^ secondary antibody. Images were acquired on an Olympus FV1000 confocal microscope. Scale bar= 5 μM.

### Sertraline inhibits uptake of the dynamin-dependent cargo transferrin

Unlike imipramine and citalopram, sertraline did not enhance Lp(a) uptake. Sertraline has established, potent off-target inhibitory effects on the mediator of endosome scission, dynamin (Otomo et al., 2008), which prevents endocytosis of the dynamin-dependent cargo transferrin (Takahashi et al., 2010). While macropinocytosis is conflictingly described as dynamin-dependent and dynamin-independent (Cao et al., 2007; Li et al., 2015), constitutive macropinocytosis (as opposed to growth factor activated macropinocytosis) is dependent on dynamin in a wide range of cell types including primary rat hepatocytes (Cao et al., 2007; Kolpak et al., 2009). Given the hepatic origin of the HepG2 cell line and the exclusion of serum (and therefore growth factors) in our experiments, constitutive macropinocytosis is the process likely active in our study. To confirm that sertraline is inhibiting dynamin-dependent endocytosis in our experimental system, we quantified uptake of the dynamin-dependent cargo transferrin in the presence of sertraline. As expected, sertraline inhibited the uptake of transferrin (Figure S3A,B). These results indicate that the effects of sertraline as a dynamin inhibitor may prevent it enhancing Lp(a) uptake as imipramine and citalopram do.

### Imipramine and serotonin upregulate PlgRKT

Anchoring of Lp(a) to the cell surface by PlgRKT is an important step in Lp(a) uptake, followed by S100A10 and annexin A2 mediated macropinocytosis (Siddiqui et al., 2023). As imipramine, citalopram and serotonin all enhanced Lp(a) binding to the cell surface and subsequent uptake, we investigated the effects of these compounds on expression of PlgRKT, S100A10 and annexin A2. We evaluated expression of each via immunofluorescence due to our inability to find antibodies that reliably detect all three proteins by Western blot. PlgRKT, S100A10 and annexin A2 antibodies were previously validated to be specific via immunofluorescence (Siddiqui et al., 2023). Imipramine and serotonin, but not citalopram, induced significant increases in PlgRKT immunofluorescence in HepG2 cells (Figure 4A-F). Citalopram and serotonin, but not imipramine, significantly upregulated expression S100A10 (Figure S4A-F), while all three compounds had no effect on annexin A2 expression (Figure S4G,H). Imipramine and serotonin most likely enhance Lp(a) uptake by increasing its PlgRKT-dependent plasma membrane anchoring. While citalopram (and serotonin) upregulated S100A10, this upregulation is unlikely to stimulate Lp(a) macropinocytosis as S100A10 requires annexin A2 to function as a heterotetrametric complex in endocytic processes (Morel and Gruenberg, 2007). Indeed, citalopram and serotonin had no effect on dextran uptake (Figure S1B, Figure 3I) supporting a lack of an effect on macropinocytosis. Citalopram therefore appears to act via a mechanism other than upregulation of plasminogen receptors.

**Figure 4:**
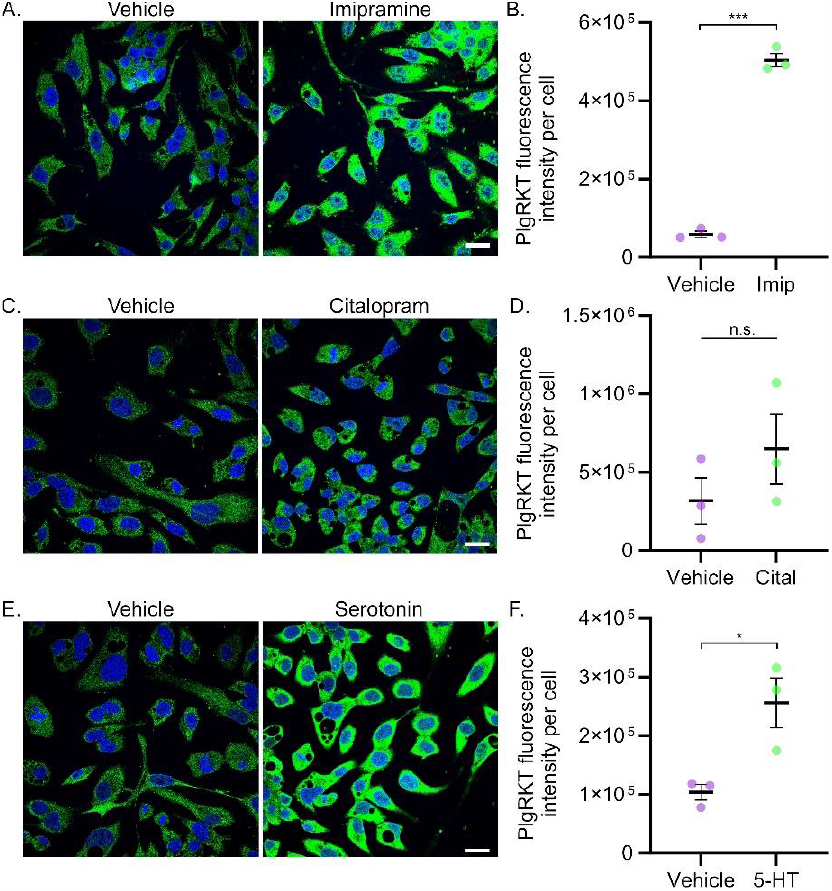
Imipramine and serotonin enhance PlgRKT expression. **(A)** Representative images of PlgRKT in HepG2 cells incubated with H_2_O vehicle or 20 μM imipramine (Imip) overnight. Cells were treated overnight, fixed with PFA, permeabilised with 0.15% triton-X100 and incubated with anti-PlgRKT which was detected with AlexaFluor^488^. **(B)** Quantification of PlgRKT fluorescence intensity in vehicle and imipramine treated cells. **(C)** Representative images of PlgRKT in HepG2 cells incubated with H_2_O vehicle or 50 μM citalopram (Cital) overnight. Cells were treated overnight, fixed with PFA, permeabilised with 0.15% triton-X100 and incubated with anti-PlgRKT which was detected with AlexaFluor^488^. **(D)** Quantification of PlgRKT fluorescence intensity in vehicle and citalopram treated cells. **(E)** Representative images of PlgRKT in HepG2 cells incubated with H_2_O vehicle or 500 μM serotonin (5-HT) overnight. Cells were treated overnight, fixed with PFA, permeabilised with 0.15% triton-X100 and incubated with anti-PlgRKT which was detected with AlexaFluor^488^. **(F)** Quantification of PlgRKT fluorescence intensity in vehicle and serotonin treated cells. n.s.= not significant, *= p<0.05, ***= p<0.001 from Students t-test. Data points represent means of independent experiments quantified from 5 fields of view of ∼20 cells per field from each independent experiment. Error bars represent standard error of the mean. Images were acquired on an Olympus FV1000/FV1200 confocal microscope. Scale bar= 20 μM.

### Imipramine and citalopram stimulate Lp(a) and apo(a) delivery into recycling endosomes

We previously reported that following internalisation, the apo(a) protein component of Lp(a) is predominantly recycled into the extracellular environment via Rab11-positive recycling endosomes (Sharma et al., 2017), as well as a consistent low-level of delivery into lysosomal compartments (Siddiqui et al., 2023). We therefore sought to establish if imipramine and citalopram induced Lp(a) delivery into recycling endosomes or lysosome compartments for recycling or degradation, respectively.

Lp(a) was added to HepG2 cells for 30 or 120 minutes following overnight imipramine or citalopram treatment, cells fixed, permeabilised and co-stained for Lp(a) and Rab11 (recycling endosomes), Rab7 (late endosomes) or LAMP1 (lysosomes). Cells were then imaged by confocal microscopy and co-localisation between Lp(a) and endosomal compartment markers quantified by object-based colocalisation using threshold-defined endosomal signals to exclude plasma membrane signal.

After 30 minutes incubation, Lp(a) co-localisation with Rab11 was significantly increased with imipramine and citalopram treatments compared to vehicle control (Figure 5A,B). No significant difference in Lp(a) and Rab11 co-localisation was observed between vehicle control and imipramine or citalopram following 120 minutes of Lp(a) incubation (Figure 5A,C). No significant difference between vehicle control and imipramine/citalopram treated cells was observed in Lp(a) co-localisation with either Rab7 (Figure S5A-C) or LAMP1 (Figure S5C-F) at any timepoint, confirming that imipramine and citalopram stimulated Lp(a) uptake drives Lp(a) delivery into recycling, not degradative, compartments in the cell.

**Figure 5:**
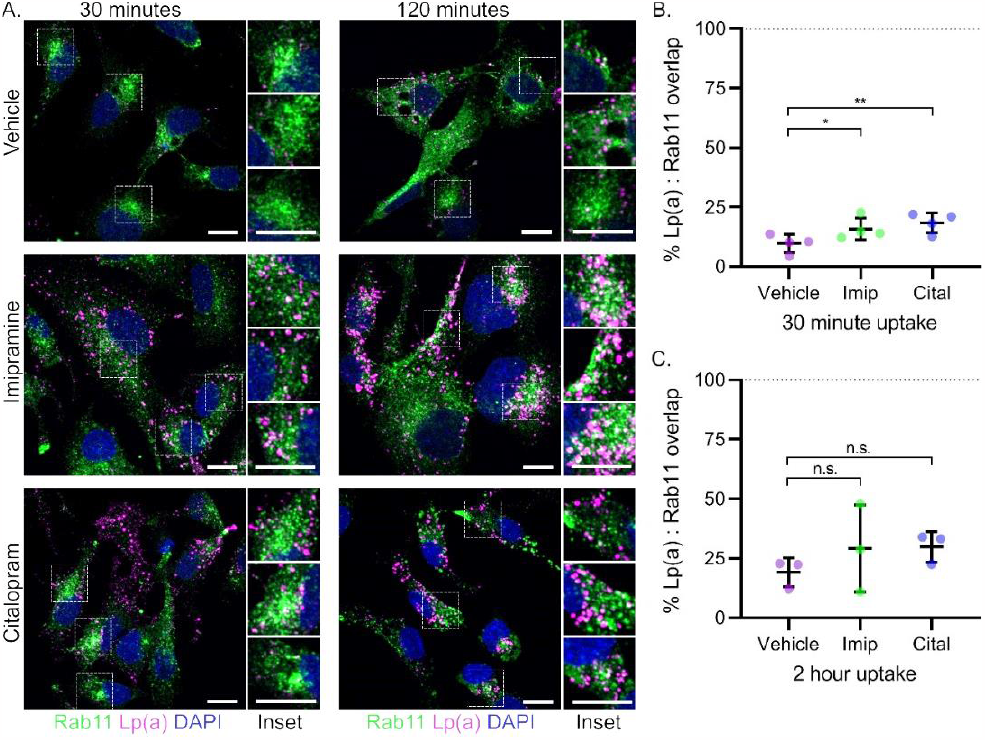
Imipramine and citalopram stimulate Lp(a) sorting into Rab11-postitive recycling endosomes. **(A)** Representative images of HepG2 cells incubated with Lp(a) for 30 minutes following H_2_O vehicle (top panel), 20 μM imipramine (middle panel) or 50 μM citalopram (bottom panel) overnight. Cells were co-stained with anti-LPA4 (magenta) and anti-Rab11 (green) and detected with AlexaFluor secondary antibodies. Quantification of percentage vesicle overlap between Lp(a) and Rab11 channels following 30 **(B)** or 120 **(C)** minutes of Lp(a) uptake. n.s.= not significant, *= p<0.05, **= p<0.01 from randomised block ANOVA comparing treated to vehicle control conditions at each timepoint. Data points represent means of independent experiments quantified from 5 fields of view of ∼20 cells per field from each independent experiment. Error bars represent standard error of the mean. Images were acquired on an Olympus FV1000/FV1200 confocal microscope. Scale bar= 5 μM.

## Discussion

Endocytosis is the central regulator of lipoprotein clearance from circulation (Zanoni et al., 2018). In our previous study we have established that Lp(a) uptake occurs via the fluid-phase endocytic mechanism, macropinocytosis (Siddiqui et al., 2023). In attempting to refine this process in our current study, we discovered that common antidepressants exert complex effects on Lp(a) uptake (Figure 6). In HepG2 liver cells, imipramine and citalopram stimulated Lp(a) uptake by increasing Lp(a) binding to the cell surface. Serotonin similarly increased Lp(a) cell surface binding and uptake *via* macropinocytosis, indicating the effects of imipramine and citalopram on Lp(a) are serotonin-dependant. Sertraline in contrast did not stimulate Lp(a) uptake, likely due to its effects as a dynamin inhibitor. Functionally, antidepressant-stimulated (imipramine and citalopram) Lp(a) uptake resulted in increased Lp(a) delivery to recycling endosomes, but not degradative compartments of the cell.

**Figure 6:**
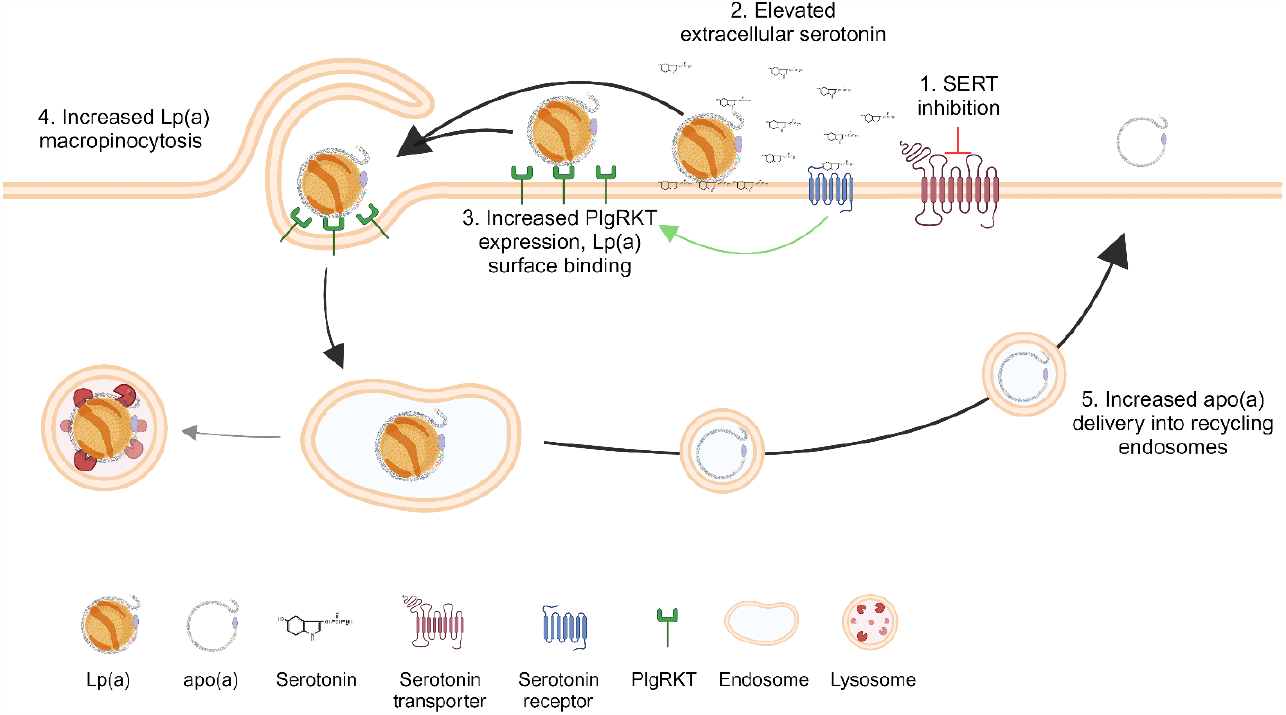
Hypothesised mechanism of serotonin and antidepressant action on Lp(a) uptake. **1)** Imipramine and citalopram inhibit the serotonin transporter, **2)** increasing extracellular serotonin levels. **3)** Serotonin increases Lp(a) binding to the cell surface by integrating into the plasma membrane and/or increasing PlgRKT expression via serotonin receptor signalling. **4)** Increased Lp(a) surface binding results in increased Lp(a) uptake by macropinocytosis. **5)** Following enhanced Lp(a) uptake, increased delivery of Lp(a) into recycling endosomes is observed, potentially leading to increased apo(a) recycling into the extracellular environment. Schematic created with BioRender.com.

Links between depression and increased atherosclerotic cardiovascular disease development have long been detected (Chávez-Castillo et al., 2018; Hare et al., 2014). Circulating Lp(a) levels have been observed to be elevated in patients with depression, an increase that correlates with unfavourable cardiovascular outcomes (Bao et al., 2021; Emanuele et al., 2006; Hamidifard et al., 2009; Hui et al., 2021). Given the contrasting effects of sertraline and citalopram on Lp(a) uptake in our cellular models, it would be of interest to determine if similar alterations in Lp(a) uptake are observed with these SSRIs *in vivo*. Interestingly, a double-blinded interventional study has shown that treatment with the SSRI paroxetine decreases circulating Lp(a) levels (Paslakis et al., 2011). We hypothesise that patients with depression and high Lp(a) levels may benefit from treatment with SSRIs that enhance Lp(a) uptake such as citalopram, which may reduce Lp(a) levels in circulation by boosting endocytosis. Conversely sertraline may not be suitable since it may increase circulating Lp(a) levels by inhibiting cellular uptake.

Serotonin has been recently discovered to be a regulator of cargoes (transferrin and islet amyloid precursor protein) binding to the plasma membrane (Dey et al., 2021). Our study extends the cargo range whose uptake is altered by serotonin, showing serotonin increased plasma membrane binding of Lp(a) to the cell surface. Dey et al., (2021) demonstrated through extensive biophysical experimentation that serotonin enhances transferrin uptake by modulating the plasma membrane biophysical properties. Here we found that serotonin can additionally act by upregulating receptors such as PlgRKT, thereby enhancing endocytosis via additional mechanisms beyond membrane modulation (Figure 6).

Questions remain from our current study that need to be addressed to better understand the effects of serotonin and antidepressants on Lp(a) uptake. Firstly, this study was entirely *in vitro* in nature, therefore *in vivo* studies would be required to determine the potential clinical relevance of our findings. Secondly, Dey et al., (2021) employed serotonin receptor inhibitors in their study to rule out receptor-dependent effects on enhanced surface binding. They found that serotonin integration into the plasma membrane induced biophysical changes that enhanced cargo plasma membrane binding and uptake. We did not employ serotonin receptor inhibitors here, therefore in addition to potential biophysical changes to the membrane induced by serotonin, serotonin receptor signalling may be in effect (Figure 6). Indeed, the upregulation of PlgRKT by imipramine and serotonin, and S100A10 by citalopram and serotonin could be due to serotonin receptor signalling. There is a well-documented interaction between S100A10 and serotonin receptor expression levels and signalling in the brain (Chen et al., 2022; Sargin et al., 2020). Our current study indicates this interplay between serotonin and S100A10 also persists in non-neuronal cell lines. While we were unable to fully dissect how citalopram enhanced Lp(a) cell surface binding and uptake, we hypothesise this may be due to a combination of the biophysical effects of serotonin observed by Dey at al., and upregulation of other components regulating Lp(a) uptake beyond the plasminogen receptors investigated here. Finally, the functional effects of imipramine and citalopram enhancement of Lp(a) delivery into recycling compartments are yet unknown. We previously found that after Lp(a) uptake into the cell, the apo(a) protein component is removed from the LDL-like component and rather than undergoing lysosomal or proteasomal degradation is re-secreted to the extracellular space (Sharma et al., 2017). Free apo(a) has been identified in circulation and urine in proteolysed forms in healthy (Edelstein et al., 1999; Mooser et al., 1996) and nephrotic syndrome patients (Doucet et al., 2000). We hypothesise that recycling may serve as a mechanism to remove the LDL-like component of Lp(a), releasing free apo(a) into circulation for proteolysis and excretion from the body.

In conclusion, our data reveal the importance of serotonin and antidepressants in modulating Lp(a) uptake via enhanced cell surface binding and macropinocytosis. Our study indicates that antidepressants, especially the SSRI citalopram, should be investigated pre-clinically and clinically for their effects on Lp(a) levels in circulation. The benefit of identifying specific SSRIs capable of reducing Lp(a) levels *in vivo* are clear, as depression has been previously linked with higher Lp(a) levels (Bao et al., 2021; Emanuele et al., 2006; Hamidifard et al., 2009; Hui et al., 2021), and depression sufferers are less likely to adhere to treatments for co-morbidities such as cardiovascular disease (Gehi et al., 2005; Goldstein et al., 2017; Grenard et al., 2011). SSRIs that reduce Lp(a) levels in depression sufferers would allow treatment of both depression and Lp(a)-induced atherosclerotic burden with a single medicine, improving patient quality of life and cardiovascular health with a better likelihood of compliance.

## Acknowledgements

The authors acknowledge Mary Blok for subject recruitment. This research was supported by project grants from the Royal Society of New Zealand (Marsden Fund 17-UOO-081) and the Health Research Council of New Zealand (18/207).

## Author contributions

ND, GMIR, HS, and GM performed the experiments. MW supervised the recruitment of Otago LPA cohort subjects; MR purified the Lp(a). GMIR and ND analysed the data. GMIR, ND and SPAM prepared the figures and wrote the manuscript. ND prepared the schematic. GR, ND and SPAM supervised the students. GR and SPAM conceived of the study.

## Methods

### Resource Availability

#### Lead contacts

Further information and requests for resources and reagents should be directed to and will be fulfilled by Professor Sally PA McCormick or Dr. Gregory MI Repdath.

#### Data and Code Availability

The raw confocal imaging data has been deposited into the FigShare repository. FIJI Macros are available upon request from Dr. Gregory Redpath.

### Experimental Model and Subject Details

#### Cell culture

Cells were grown at 37°C in a 5% CO_2_ humidified incubator. HepG2 cells (ATCC, CVCL-2205) and HEK293 cells (ATCC, CVCL_0045) were cultured in DMEM with 10% FBS, 100 u/ml penicillin, 100μg/ml streptomycin, and 0.25 mg/mL amphotericin B.

#### Lp(a) enrichment from patient samples

Lp(a) was isolated from plasma samples from healthy individuals from the Otago LPA cohort (Morgan et al., 2020) with Lp(a) levels >50 mg/dL (equivalent to 100 nmol/L using a conversion factor of 2). Individual samples used for isolation had apo(a) isoform sizes ranging from 17-20 with a predominant apo(a) isoform size of 19. 100 μg total protein concentration of enriched Lp(a) (equivalent to 88–178 nM) was used per experimental condition. Lp(a) protein levels were quantified using the Qubit Protein Assay Kit (Thermo Fisher Scientific) and characterised for enrichment as described by Siddiqui et al., (2023).

#### Ethical Approval

Ethical approval for the samples used in this study was granted by the Lower Regional South Ethics Committee (LRS/12/01/005).

### Method Details

#### Drug treatments

For serotonin, imipramine and citalopram treatments, cells were treated overnight prior to the experiment. Two hours before addition of Lp(a)/apo(a), cells were transferred into serum-free media and new serotonin/imipramine/citalopram added. For sertraline treatment in HepG2 cells, cells were transferred into serum-free media two hours prior to Lp(a)/apo(a) addition. Two hours before addition of Lp(a)/apo(a), cells were transferred into serum-free media and new sertraline added.

#### Immunofluorescence staining

Cells were fixed with 4% paraformaldehyde for 15 minutes at 37°C in a humidified incubator. Cells were then washed with PBS and blocked with 3% goat serum. The antibodies used in this study were: anti-apolipoprotein(a) antibody, clone LPA4 (Millipore), rabbit monoclonal anti-Lp(a) antibody (Abcam), goat polyclonal anti-Lp(a) antibody (WAKO), rabbit monoclonal anti-Rab7 (Abcam), rabbit anti-LAMP1 (Cell Signaling Technologies) and rabbit anti-Rab11 (Cell Signaling Technologies). The secondary antibodies include goat anti-rabbit Alexa Fluor 594, anti-mouse Alexa Fluor 488, and anti-rabbit Alexa Fluor 594 (ThermoFisher Scientific, 1:1000). Slides were mounted with ProLong™ antifade mounting medium (Invitrogen).

#### Confocal imaging

All coverslips were imaged on an Olympus FV1000/1200/3000 confocal microscope using 405, 488 and 561 nm laser lines with a 60x, NA 1.4 oil immersion lens. Within each experiment, images from each coverslip were captured using identical laser power, gain, offset and PMT (FV1000/1200)/GaAsP(FV3000) detector voltage. The maximum field of view was captured at a pixel size of ∼125-135 nm.

#### Apo(a) expression and concentration

Apo(a)-mScarlet was transfected into HEK293 cells using Lipofectamine 3000. 4 days post transfection, cell media was collected, cooled and centrifuged at 3000g to pellet cellular debris. Media was then transferred into a 20 mL, 100 kDa MW cutoff centrifuge tube (Thermo Fisher Scientific) and centrifuged at 3000g at 4°C until media volume was 5x concentrated. Concentrated media was frozen until used.

### Quantification and Statistical Analysis

#### Image Quantification

Images were quantified using FIJI(Schindelin et al., 2012). Custom macros were created for endosome identification and overlap analysis. For endosome identification, a single intensity-based threshold per experiment was used to identify Lp(a)/apo(a) endosomes and nuclei. For overlap analysis, one intensity-based threshold was used to identify Lp(a)/apo(a) endosomes, and another used to identify Rab7/11/LAMP1 compartments. The same threshold was used for each image in a single experiment. The threshold was used to generate a mask for each channel, and the Lp(a)/apo(a) mask subtracted from the inverted Rab7/11/LAMP1 mask to generate an overlap mask of Lp(a)/apo(a) present in the Rab7/11/LAMP1 compartment. The number of Lp(a)/apo(a) endosomes in the overlap mask was quantified and divided by the total number of Lp(a)/apo(a) endosomes to determine the percentage of total Lp(a)/apo(a) endosomes overlapping with Rab7/11/LAMP1 compartments.

#### Data Representation and Statistical analysis

Data are expressed as means of independent experiments +/- SEM, excepting Figure S3 in which data points represent individual fields of view. Each independent experiment consists of 5 fields of view containing approximately 20 cells per field. All datasets were tested for normality and the corresponding parametric or non-parametric tests used based upon this. Statistical analyses were performed with Prism Software version 8 (GraphPad Software

### Resources Table

| REAGENT or RESOURCE | SOURCE | RRID IDENTIFIER |
| --- | --- | --- |
| **Antibodies** |  |  |
| Rabbit anti-LAMP1 | Cell Signaling Technology | Cat# 9091; RRID: AB_2687579 |
| Anti-Apolipoprotein(a) antibody, clone LPA4 | Millipore | Cat#MABS1284 |
| Rabbit monoclonal anti-Rab7 [EPR7589] | Abcam | Cat#ab137029; RRID: AB_2629474 |
| Rabbit anti-Rab11 | Cell Signaling Technology | Cat# 5589; RRID: AB_10693925 |
| Rabbit monoclonal anti-Lp(a) antibody | Abcam | Cat# ab208184 |
| Anti-PLGRKT polyclonal antibody | Abcam | Cat# ab237525 |
| Anti-annexin A2 monoclonal antibody | Cell Signaling Technology | Cat#8235; RRID: RRID:AB_11129437 |
| Anti-S100A10 monoclonal antibody | Cell Signaling Technology | Cat#5529; RRID: AB_10834810 |
| Donkey anti-Mouse IgG (H+L) Highly Cross-Adsorbed Secondary Antibody, Alexa Fluor 488 | Thermo Fisher Scientific | Cat# A-21202, RRID:AB_141607 |
| Donkey anti-Mouse IgG (H+L) Highly Cross-Adsorbed Secondary Antibody, Alexa Fluor 594 | Thermo Fisher Scientific | Cat# A-21203, RRID:AB_141633 |
| Donkey anti-Rabbit IgG (H+L) Highly Cross-Adsorbed Secondary Antibody, Alexa Fluor 488 | Thermo Fisher Scientific | Cat# A-21206, RRID:AB_2535792 |
| Donkey anti-Rabbit IgG (H+L) Highly Cross-Adsorbed Secondary Antibody, Alexa Fluor 594 | Thermo Fisher Scientific | Cat# A-21207, RRID:AB_141637 |
| **Biological Samples** |  |  |
| Purified human Lp(a) |  |  |
| **Chemicals, Peptides, and Recombinant Proteins** |  |  |
| Phosphate buffered saline, pH 7.4 | Gibco | Cat#10010023 |
| DMEM, high glucose | Gibco | Cat#11965092 |
| Fetal Bovine Serum, qualified, New Zealand | Gibco | Cat#10091155 |
| Penicillin-Streptomycin (10,000 U/mL) | Gibco | Cat#15140122 |
| Amphotericin B | Gibco | Cat#15290018 |
| TrypLE™ Express Enzyme (1X), no phenol red | Gibco | Cat#12604021 |
| Dextran, Tetramethylrhodamine, 70,000 MW, Lysine Fixable | Invitrogen | Cat#D1818 |
| ProLong™ antifade mounting reagent | Invitrogen | Cat#P36934 |
| Triton-X100 | Bio-Rad Laboratories | Cat#1610407 |
| Imipramine hydrochloride | Sigma-Aldrich | Cat#I7379 |
| Sertraline tablets 50 mg as hydrochloride | Teva Pharmaceutical Industries Limited |  |
| Citalopram tablets 20 mg as hydrobromide | PSM Healthcare Limited |  |
| Serotonin hydrochloride (5-Hydroxytryptamine) | Abcam | Cat#ab120528 |
| Paraformaldehyde | Sigma-Aldrich | Cat#P6148 |
| Goat Serum, New Zealand origin | Sigma-Aldrich | Cat#16210064 |
| Qubit Protein Assay Kit | Thermo Fisher Scientific | Cat#Q33211 |
| Pierce Protein Concentrator PES, 100K MWCO, 5-20 mL | Thermo Fisher Scientific | Cat#88532 |
| Lipofectamine 3000 | Life Technologies | Cat#L3000015 |
| **Deposited Data** |  |  |
| Figshare |  |  |
| **Experimental Models: Cell Lines** |  |  |
| HepG2 | ATCC | ATCC Cat# HB-8065, RRID:CVCL_0027 |
| HEK293 | ATCC | ATCC Cat# CRL-1573, RRID:CVCL_0045 |
| **Recombinant DNA** |  |  |
| Lipoprotein a (LPA) (NM_005577) Human Tagged ORF Clone | Origene | CAT#RG212070 |
| **Software and Algorithms** |  |  |
| Prism | Graphpad | RRID:SCR_002798 |
| FIJI |  | RRID:SCR_002285 |
| Fluoview FV-10 ASW | Olympus | RRID:SCR_014215 |
| BioRender |  | RRID:SCR_01836 |

