## Supplementary Figures for "Antidepressants stimulate lipoprotein(a) macropinocytosis via serotonin-enhanced cell surface binding"

**Supplementary Information**

**Figure S1: Antidepressants do not enhance dextran uptake in liver cells. (A)** Representative images of HepG2 cells incubated with 70kDa dextran-TRITC for 1 hour following H_2_O vehicle (left to right), 20 µM imipramine (overnight), 50 µM citalopram (overnight), and 20 µM sertraline (2 hours) treatment. **(B)** Quantification of 70 kDa dextran-TRITC vesicles following overnight treatment with imipramine (imip) and citalopram (cital) compared to vehicle control (veh). **(C)** Quantification of 70 kDa dextran-TRITC vesicles following 2 hour treatment with sertraline (sert) compared to vehicle control (Veh). n.s.= not significant from a randomised-block ANOVA with Friedman test for multiple comparisons **(B)**, **=p<0.01 from a Student’s T-test **(C)**. Data points represent means of independent experiments quantified from 5 fields of view of ~20 cells per field from each independent experiment. Error bars represent standard error of the mean. Dextran was detected directly using an Olympus FV1000/FV1200 confocal microscope. Scale bar= 5 µM.

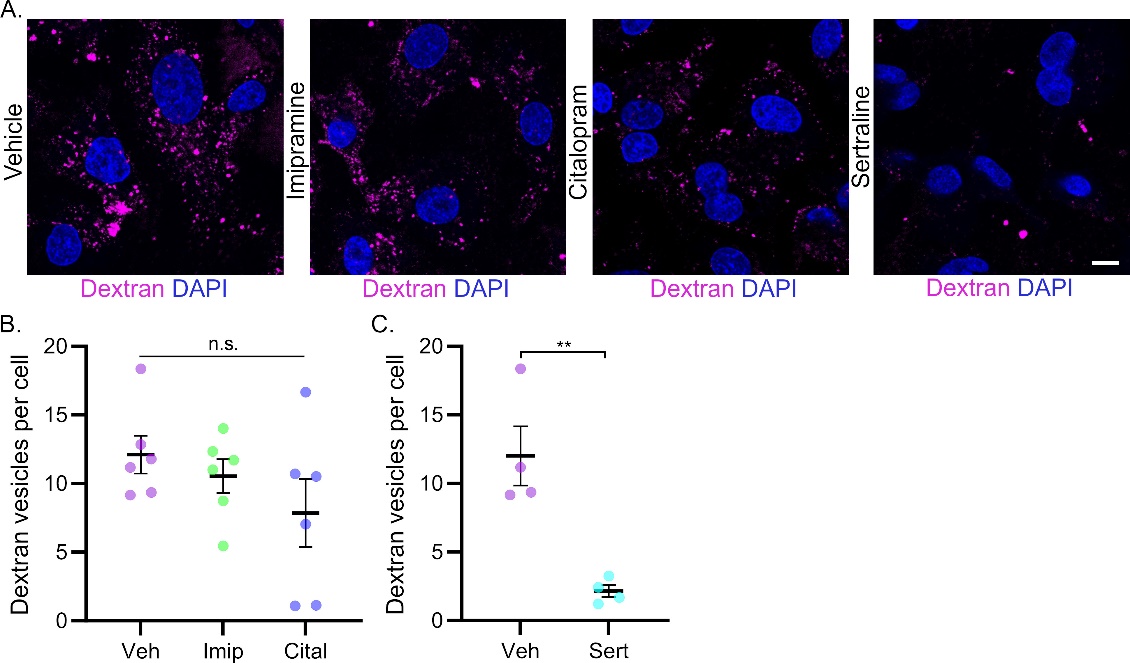

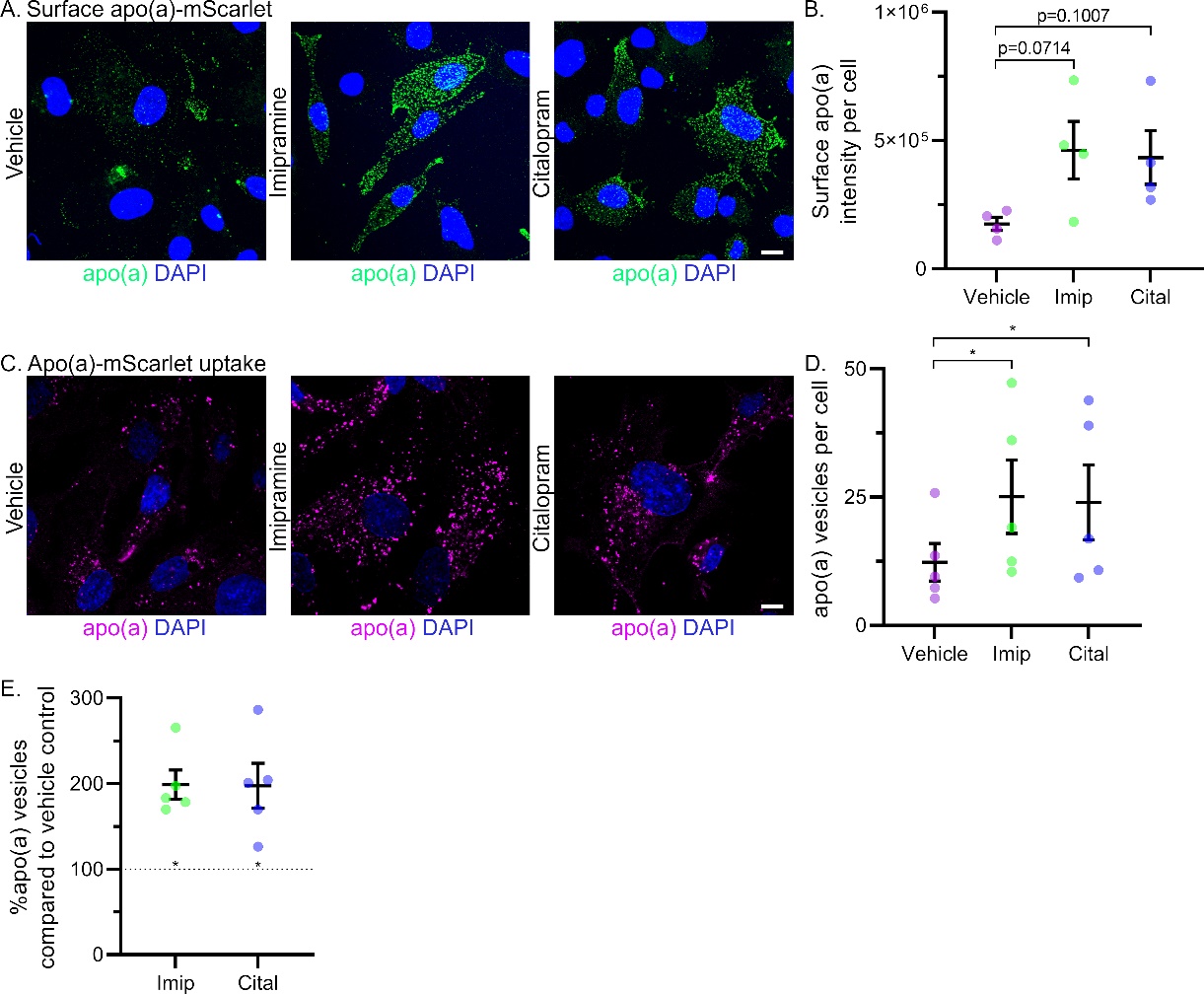

**Figure S2: Antidepressants increase apo(a) surface binding and uptake in liver cells. (A)** Representative images of maximum z-projections of surface-bound apo(a)-mScarlet following H_2_O vehicle, 20 µM imipramine or 50 µM citalopram treatment overnight. **(B)** Surface apo(a) signal per cell in vehicle, imipramine and citalopram treated cells. **(C)** Representative images of apo(a)-mScarlet following H_2_O vehicle, 20 µM imipramine or 50 µM citalopram treatment overnight and incubation with apo(a)-mScarlet for 120 minutes. **(D)** Apo(a)-mScarlet vesicles detected following 2-hour apo(a) incubation in HepG2 cells treated with H_2_O vehicle, imipramine or citalopram overnight. Apo(a)-mScarlet was detected using the LPA4 antibody and anti-mouse^488^ **(A)** or the LPA4 antibody with anti-mouse^594^. Images were acquired on an Olympus FV1000/FV1200 confocal microscope. Data points present means of independent experiments. Error bars represent standard error of the mean. P values or *=p<0.05 from randomised block ANOVA.

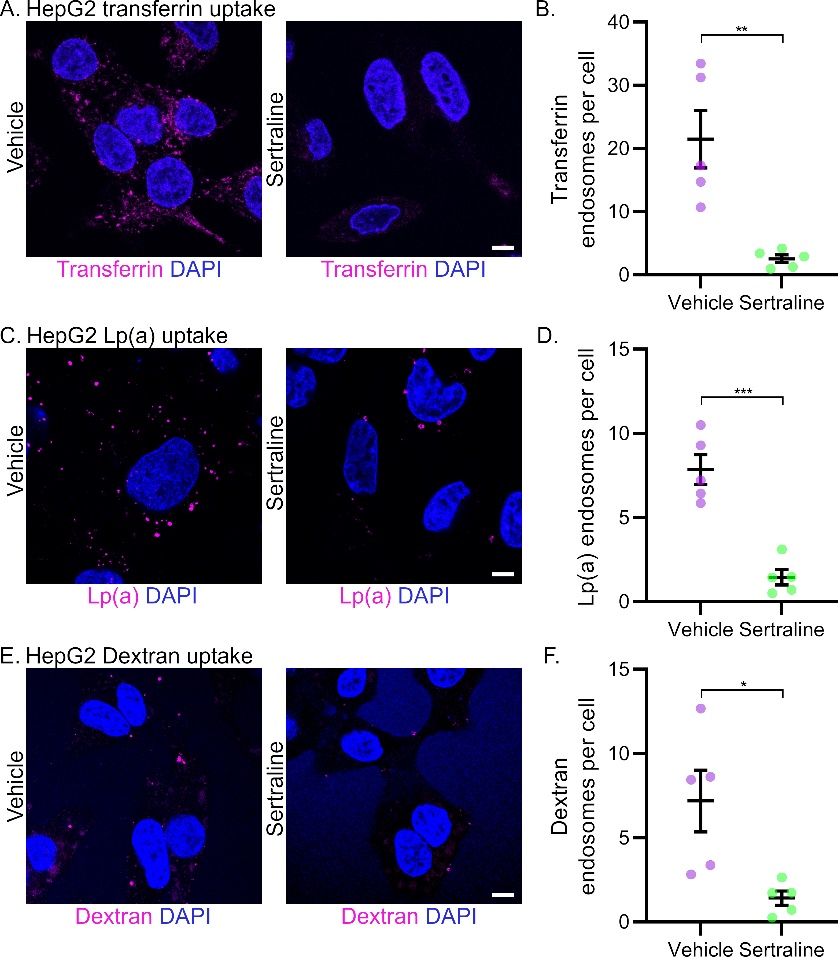

**Figure S3: Sertraline inhibits transferrin uptake. (A)** Representative images of HepG2 cells treated vehicle (left) or sertraline (right) for 2 hours, followed by transferrin-594 for 1 hour. **(B)** Transferrin vesicles detected following 1 hour transferrin incubation in HepG2 cells +/- sertraline. Data points represent mean vesicles from a single field of view, error bars represent mean +/- SD from one independent experiment. **=p<0.01 from unpaired Student’s T-test. Images were acquired on an Olympus FV3000 confocal microscope. Scale bar= 5 µM.

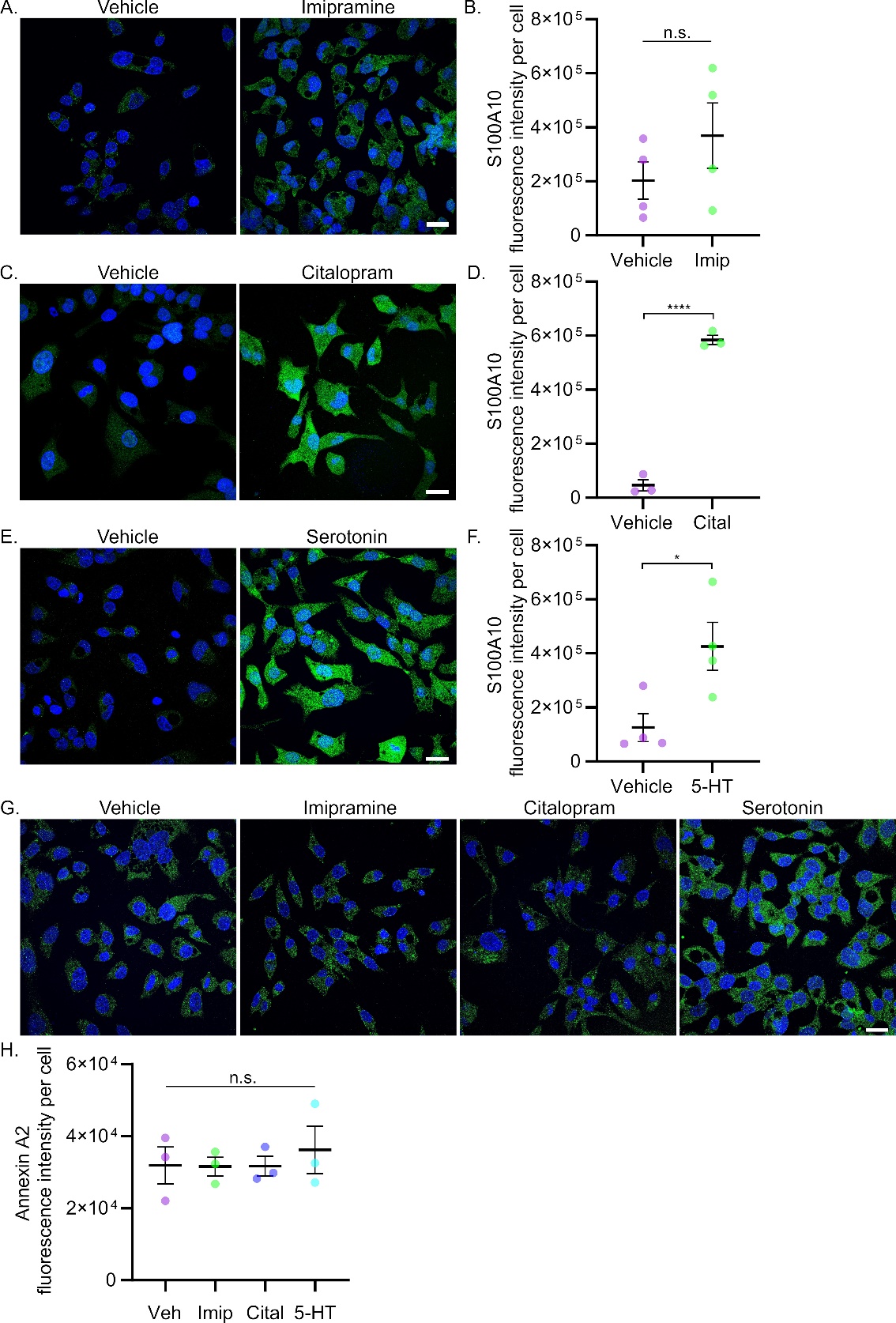

**Figure S4: Citalopram and serotonin upregulate S100A10, but not Annexin A2. (A)** Representative images of S100A10 expression in HepG2 cells treated with H_2_O vehicle or 20 µM imipramine overnight. **(B)** Quantification of S100A10 intensity per cell following vehicle or imipramine treatment. **(C)** Representative images of S100A10 expression in HepG2 cells treated with H_2_O vehicle or 50 µM citalopram overnight. **(D)** Quantification of S100A10 intensity per cell following vehicle or citalopram treatment. **(E)** Representative images of S100A10 expression in HepG2 cells treated with H_2_O vehicle or 500 µM serotonin overnight. **(F)** Quantification of S100A10 intensity per cell following vehicle or serotonin treatment. **(G)** Representative images of Annexin A2 expression in HepG2 cells treated with H_2_O vehicle, 20 µM imipramine, 50 µM citalopram or 500 µM serotonin overnight. **(H)** Quantification of S100A10 intensity per cell following vehicle (veh), imipramine (imip), citalopram (cital) or serotonin (5-HT) treatment. n.s= not significant, *= p<0.05, ****= p<0.0001 from Student’s T-test **(B,D,F)**. n.s.= not significant from randomised block ANOVA comparing treated to vehicle control **(H)**. Data points represent means of independent experiments quantified from 5 fields of view of ~20 cells per field from each independent experiment. Error bars represent standard error of the mean. Images were acquired on an Olympus FV1000/FV1200 confocal microscope. Scale bar= 5 µM.

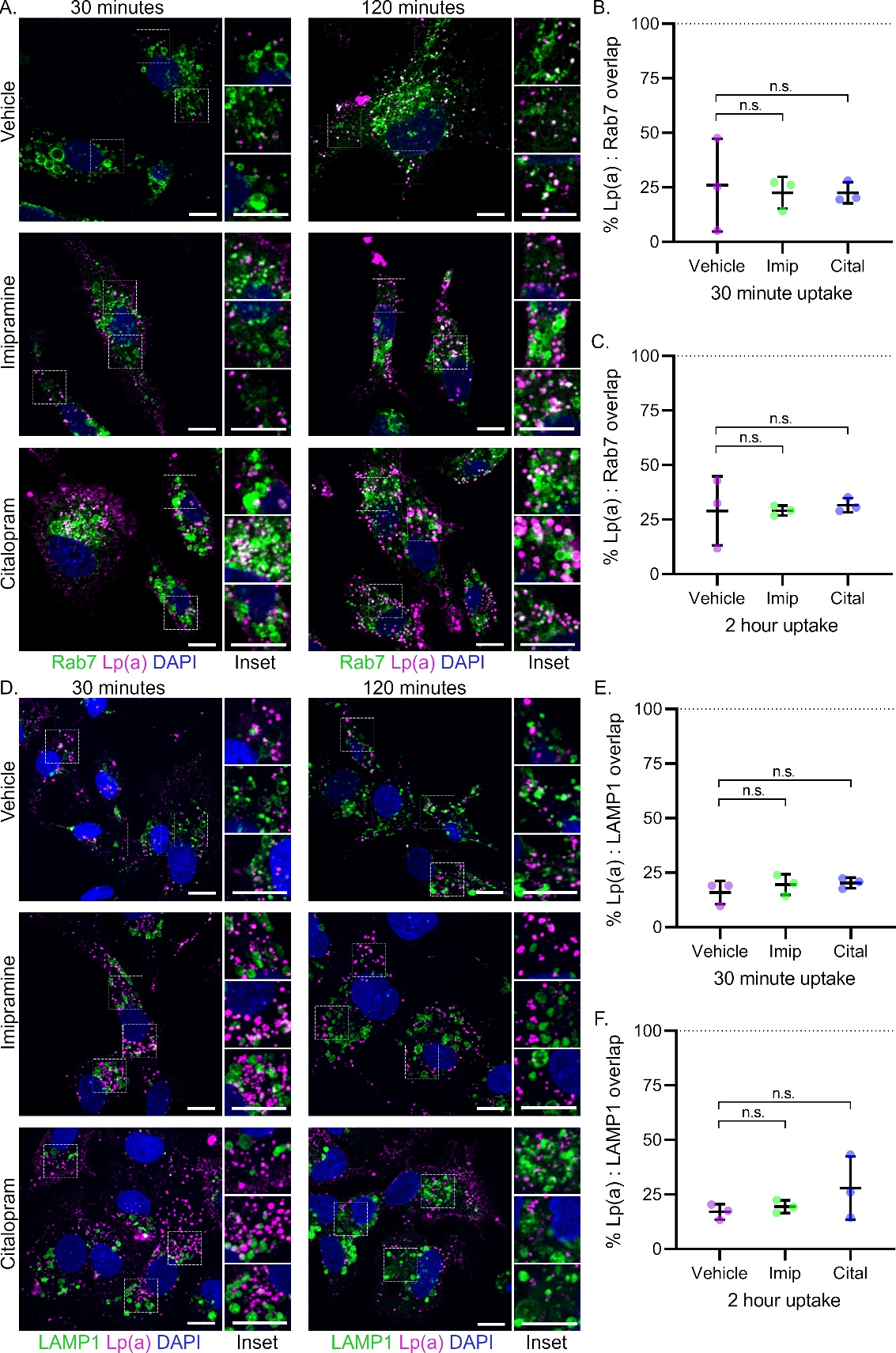

**Figure S5: Imipramine and citalopram do not change Lp(a) delivery to late endosomal and lysosomal compartments. (A)** Representative images of HepG2 cells incubated with Lp(a) for 30- or 120-minutes following H_2_O vehicle (top panel), 20 µM imipramine (middle panel) or 50 µM citalopram (bottom panel) overnight. Cells were co-stained with anti-LPA4 (magenta) and anti-Rab7 (green) and detected with AlexaFluor secondary antibodies. Quantification of percentage vesicle overlap between Lp(a) and Rab7 channels following 30 **(B)** or 120 **(C)** minutes of Lp(a) uptake. **(D)** Representative images of HepG2 cells incubated with Lp(a) for 30- or 120-minutes following H_2_O vehicle (top panel), 20 µM imipramine (middle panel) or 50 µM citalopram (bottom panel) overnight. Cells were co-stained with anti-LPA4 (magenta) and anti-LAMP1 (green) and detected with AlexaFluor secondary antibodies. Quantification of percentage vesicle overlap between Lp(a) and LAMP1 channels following 30 **(E)** or 120 **(F)** minutes of Lp(a) uptake n.s.= not significant, *= p<0.05, **= p<0.01 from randomised block ANOVA comparing treated to vehicle control conditions at each timepoint. Data points represent means of independent experiments quantified from 5 fields of view of ~20 cells per field from each independent experiment. Error bars represent standard error of the mean. Images were acquired on an Olympus FV1000/FV1200 confocal microscope. Scale bar= 5 µM.
